# Aging promotes inflammation and steatosis in alcohol-associated liver disease in mice

**DOI:** 10.64898/2026.06.19.730442

**Authors:** Sha Neisha Williams, Xiaowen Ma, Xiaojuan Chao, Hangfei Xu, Wanqing Liu, Hong-Min Ni, Wen-Xing Ding

**Author notes:** Correspondence to: Wen-Xing Ding, Ph.D., Department of Pharmacology, Toxicology and Therapeutics, University of Kansas Medical Center, MS 1018, 3901 Rainbow Blvd., Kansas City, Kansas 66160.

## Abstract

**Background and Aims:** As the older population (aged 65 years and older) continues to expand and more people drink alcohol, aging has been linked to the development of alcohol-associated liver disease (ALD) and to worse disease outcomes. The aim of this study was to explore the mechanisms by which advanced age and alcohol exacerbate alcohol-induced liver injury.

**Methods:** Two-to-three-month-old and twenty-to-twenty-two-month-old male C57BL/6N mice were subjected to chronic-on-binge alcohol feeding (the Gao-binge model). Serum alanine aminotransferase, aspartate aminotransferase, and triglyceride content were determined using biochemical assays. The levels of lipogenesis, fatty acid-metabolizing proteins, inflammatory markers, mitochondrial and autophagy-related proteins, and senescence-associated proteins were determined by immunoblotting, immunohistochemistry, and real-time quantitative polymerase chain reaction (RT-qPCR). Liver tissues were also subjected to RNA sequencing and metabolomics analyses. Proteomics analysis was performed on serum samples. Tail-vein adenovirus-TFEB was injected to overexpress hepatic TFEB in 22-month-old C57BL/6N male mice, followed by Gao-binge alcohol feeding.

**Results:** Hepatic triglyceride content was significantly increased in aged, alcohol-fed mice, whereas serum ALT and AST levels remained relatively similar between alcohol-fed young and aged mice. Gao-binge alcohol increased the hepatic levels of diacylglycerol and acyl-carnitine species in both aged and young livers. RNA sequencing, proteomic analysis, and serum cytokine array analysis showed that inflammatory cytokines, including Ccr2, Cxcl1, and CCL6, and pro-inflammatory antibody fragments were increased in aged, alcohol-fed mice. Increased gene and protein expression of the senescent markers p21 and p27, along with increased senescent-associated (SA) β-galactosidase activity in ethanol-fed aged mice compared to young mice. Gene and protein expression of TFEB was downregulated in ethanol-fed young and aged animals, along with decreased levels of lysosomal ATPases and hepatic dipeptide content. Overexpression of TFEB in the livers of aged, Gao-binge-fed mice was associated with reduced Ly6G-positive cells, reduced protein levels of the innate immune mediators cGAS, IRF-7, IRF3, and NLRP3, and reduced caspase-1 activity as well as serum ALT levels.

**Conclusions:** Our findings indicate that advanced age perpetuates the detrimental effects of excessive alcohol consumption on various homeostatic processes and promotes steatosis and inflammation in the liver. Modulations in hepatic TFEB could be effective in mitigating pro-inflammatory signaling that occurs due to the synergistic effect of both heavy alcohol consumption and advanced age.

## 1. Introduction

In 2025, the Substance Abuse and Mental Health Services Administration (SAMHSA) collected data on alcohol use from 2024 in the United States. Analysis of the results from this survey revealed that more than 60% of the adult population (18 years and older) had consumed alcohol within the past year. Though the possible beneficial effects of moderate drinking are debatable, many studies and reviews have been published to confirm the detrimental consequences of excessive alcohol consumption (Williams, Manley et al. 2014, Nagy, Ding et al. 2016, Mackowiak, Fu et al. 2024). The harmful effects associated with heavy drinking encompass a wide range of disease phenotypes that, at first, are highlighted by the hepatic accumulation of lipids, or steatosis, which progress into more severe conditions such as steatohepatitis, cirrhosis of the liver, and even liver cancer (Massey and Arteel 2012, Ceni, Mello et al. 2014, Mackowiak, Fu et al. 2024). The onset of these various conditions results from the impairment of several homeostatic processes, notably autophagy, lipid metabolism, cellular redox regulation, endoplasmic reticulum (ER) stress, and mitochondrial quality control resulting in increased oxidative stress, megamitochondria and Mallory-Denk body formation, ductular reactions and increased innate immune response and infiltration of inflammatory cells in the liver (Chao, Wang et al. 2023, Ma, Ni et al. 2024, Mackowiak, Fu et al. 2024, Hinz, Qian et al. 2025, Ma, Niu et al. 2026). Collectively, this disease spectrum is known as alcohol-associated liver disease (ALD) and is one of the leading causes of alcohol-associated mortality, with a prevalence rate of around 4.7%, globally (Mackowiak, Fu et al. 2024, Narro, Diaz et al. 2024).

In the SAMHSA survey highlighted above, alcohol use and misuse amongst various age groups from the previous year were reported as well. Among those groups assessed, ∼15% of survey participants from the older adult population (>65 years of age) reported being binge and/or heavy drinkers. Though this number may seem low in comparison to binge drinking rates in other age groups, the number of adults in the older adult population is steadily increasing (U.S. Census Bureau, 2023), potentially raising the percentage of binge drinkers in this age group adding to the economic burden associated with ALD (Julien, Ayer et al. 2024). As the liver ages, structural, cellular, and molecular changes occur that increase susceptibility to developing chronic liver diseases (Kim, Kisseleva et al. 2015, Hunt, Kang et al. 2019, Le Couteur, Ngu et al. 2025, Williams and Ding 2025). These events include diminished drug absorption and clearance, increased production and circulation of senescent-associated molecules, loss of efficient cellular clean-up mechanisms, persistent low-grade inflammation, and overall changes to liver structure (Williams and Ding 2025).

Another function integral to maintaining liver health, which is affected by aging, is autophagy (Williams and Ding 2025). In general, autophagy functions in the maintenance of liver homeostasis by breaking down damaged organelles and by regulating hepatic metabolism through the degradation of lipids, proteins, and carbohydrates to provide energy products for hepatic cells to utilize during periods of cellular stress such as aging, starvation, or even acute ethanol consumption. Additionally, autophagy is critical in mitigating inflammatory signaling as it aids in the removal of potent inflammatory activators as well as reduces cytokine production and inflammasome activation (Ueno and Komatsu 2017, Qian, Chao et al. 2021). As a regulator of lysosome-mediated autophagy, Transcription Factor-EB (TFEB), was found to have an age-dependent decline in protein content and activity in organs like the brain and proximal tubules in animal studies (Wang, Muthu Karuppan et al. 2021, Chen, Hinz et al. 2024, Nakamura, Yamamoto et al. 2024). Furthermore, decreased hepatic TFEB and function was associated with a relatively short period of ethanol exposure in mice, resulting in insufficient hepatic autophagy (Chao, Wang et al. 2018).

Currently, the dynamic changes that occur in alcohol-induced liver injury in the aged liver are still not thoroughly understood. The aim of the present study is to further evaluate the synergistic impairment of the liver induced by old age and chronic-on-binge alcohol consumption, as well as to determine whether boosting TFEB-mediated autophagy would improve age-associated ALD in mice.

## 2. Methods and Materials

### 2.1 Gao-binge alcohol model

Two-to-three-month and twenty-to-twenty-two-month-old male C57BL/6N mice (Jackson Laboratories, NIA) were subjected to the Gao-binge model for alcohol feeding as described previously (Bertola, Mathews et al. 2013). Briefly, mice were administered the Lieber-DeCarli control diet for 5 days to acclimate to the diet. On the sixth day, mice were randomly divided and either maintained on a pair-fed control diet or given an ethanol diet (plus 5% ethanol) for 10 days. After the feeding period, mice were removed from the liquid diets and administered either an ethanol gavage (5 g/kg) or a maltose-dextrin gavage (9 g/kg). Mice were allowed to recover for 8 hours, then euthanized for the collection of serum and liver tissue.

### 2.2 Biochemical assays

Activity assays for both alanine transaminase (ALT, Pointe Scientific, A7526-625, Canton, MI) and aspartate aminotransferase (AST, Pointe Scientific, A7561150, Canton, MI) were used to assess liver injury in serum samples as previously described (Qian, H 2023). Serum triglycerides and serum ethanol concentrations were measured using colorimetric kits as previously described (Qian, Chao et al. 2023).

### 2.3 Cytokine Array

The Proteome Profiler Mouse XL Cytokine Array Kit (R&D Systems, ARY028, Wiesbaden, Germany) was used to determine the serum cytokine profile. Mouse serum from 4 animals per group (10μL per mouse) were pooled and processed according to the manufacturer’s instructions. Densitometry of the blots was quantified using Fiji for ImageJ software (NIH).

### 2.4 Western blot analysis

Total liver lysates were homogenized using radioimmunoprecipitation assay (RIPA) buffer (1% NP40, 0.5% sodium deoxycholate, 0.1% sodium dodecyl (lauryl) sulfate) supplemented with protease inhibitor cocktail. Protein concentration was measured using the BCA Protein Assay (Thermo Scientific, 23227, Rockford, IL). 30 µg of protein from liver lysate was used for electrophoretic separation on 10 or 12% SDS-PAGE gel and transferred to a PVDF membrane. Next, membranes were blocked for 1 hour at room temperature in 5% blocking solution containing evaporated milk and TBST (Tris-buffered Saline plus 0.1% Tween-20) and probed with the appropriate primary antibodies (Table 1) overnight at 4°C. Membranes were then incubated with secondary antibodies (Table 1) for 1 hour on the next day. After incubation in secondary antibodies, membranes were washed in TBST and were developed for protein visualization using SuperSignal West Pico Plus chemiluminescent substrate (Thermo Scientific, 34578, Rockford, IL). The membranes were imaged using Li-Cor Odessey Fc imaging system and densitometry analysis was performed with Fiji for ImageJ software (NIH).

**Table 1:** List of Antibodies used for western blots and immunohistochemistry.

| Antibody | Company | Catalogue Number |
| --- | --- | --- |
| ADH | Santa Cruz | Sc133207 |
| ALDH2 | Proteintech | 15310-1-AP |
| ALT | Abcam | ab137531 |
| β-Actin | Sigma | A5441 |
| CYP2E1 | Abcam | ab19140 |
| cGAS | Cell Signaling | 31659 |
| F4/80 | Invitrogen | 14-4801 |
| GAPDH | Cell Signaling | 2118 |
| IRF-3 | Cell Signaling | 4302 |
| IRF-7 | Cell Signaling | 72073 |
| LAMP1 | Developmental Studies Hybridoma Bank (DHSB) | 1D4B |
| LC3 | N/A | N/A |
| Ly6G | Cell Signaling | 87048S |
| NLRP3 | Cell Signaling | 15101S |
| p21 | Santa Cruz | sc-6246 |
| p27 | Neomarkers | DCS-72.F6 |
| p53 | Cell Signaling | cs32532S |
| p62 | Abnova | H00008878-M01 |
| TFEB | Bethyl | A303-673A |
| VATP6v1A1 | Proteintech | 17115-1-AP |
| VATP6v1B2 | Proteintech | 15097-1-AP |
| HRP-Goat Anti-Mouse IgG | Jackson | 115-035-146 |
| HRP-Goat Anti-Rabbit IgG | Jackson | 111-035-144 |
| HRP-Goat Anti-Rat IgG | Jackson | 112-035-143 |
| HRP Horse Anti-Rabbit IgG | Vector | MP7401 |
| HRP Horse Anti-Rat IgG | Vector | MP-7444-15 |

### 2.5 Caspase-1 activity assay

Total liver lysates made for western blot analysis were used to measure caspase-1 activity. Briefly, 20μg of total liver lysate for each sample and 2μL of caspase-1 substrate Z-YVAD-AFC (Enzo Life Sciences Inc, ALX-260-035, Farmingdale, NY) was added to a 96-well flat bottom plate (Thermo Scientific, 3296, Rockford, IL). Next, 200μL caspase buffer pH 7.2 (20mM PIPES, 100mM NaCl, 1mM EDTA (0.5M pH 8.0), 0.1% CHAPS, 10% Sucrose) was added to each well and the plate was then incubated at 37°C overnight. Fluorescence was read at 400/505nm at 50% gain using TECAN inifinite M200 Pro plate reader (Tecan, Switzerland).

### 2.6 Histology and immunohistochemistry

Paraffin-embedded liver sections (5µm thick) were stained using hematoxylin and eosin for visualization of liver histology. For immunohistochemistry, liver sections were deparaffinized followed by 3% hydrogen peroxide treatment for quenching. Tissue sections were treated with citrate buffer, pH 6.0, for heat-induced antigen retrieval. After antigen retrieval, liver sections were blocked using UltraVision Protein Block (Epredia, TA-125-PBQ, Kalamazoo, MI) to prevent non-specific background staining. Post-blocking, slides were incubated with primary antibody overnight at 4°C and with the appropriate secondary antibody at 37°C for 1 hour the next day. Positive staining was developed using ImmPACT NovaRED HRP substrate (Vector Labs, SK-4805, Newark, CA) and counterstained with hematoxylin for nuclear staining. Tissue sections were visualized at 20X magnification for positively stained areas.

### 2.7 Oil-Red-O staining

Oil-Red-O staining was performed using cryo-sectioned liver tissue as previously described (Chao, Wang et al. 2023). Briefly, livers were immersed in 4% paraformaldehyde for overnight preservation at 4°C then placed into 20% sucrose at the same temperature overnight the next day. On the following day, livers were embedded in Tissue-Tek OCT compound. Liver sections (6μm thick) were stained with Oil-Red-O in 60% isopropanol and incubated at 37°C for 15 minutes. Liver sections were then washed with 60% isopropanol followed by three washes with water, stained with hematoxylin, and washed with water as a final step.

### 2.8 Hepatic RNA isolation and reverse transcription

Frozen liver tissue from each treatment group was homogenized and used to extract RNA following the TRIzol Reagent protocol (Invitrogen, 15596018, Carlsbad, CA). Briefly, samples were mixed with chloroform and centrifuged. After centrifugation, the aqueous phase was collected and used for RNA precipitation. Samples were then mixed with 100% isopropanol for precipitation, followed by high-speed centrifugation. The pellet was washed with 75% ethanol, centrifuged, and resuspended in RNase-free water. For quantitative real-time polymerase chain reaction (rt-PCR), purified RNA was used to synthesize first-strand cDNA by reverse transcription according to the Maxima H Minus First Strand cDNA synthesis kit protocol (Thermo Scientific, K1652, Rockford, IL). Briefly, template RNA (3 µg) was mixed with Oligo(dT)18, dNTP mix, 5x RT buffer, Ribolock RNAse inhibitor, Maxima H Minus reverse transcriptase, and nuclease-free water, then amplified using a thermocycler to generate cDNA.

For RNA sequencing analysis, RNA samples were quality-checked using a Qubit assay for RNA quantification and Agilent TapeStation gel analysis for RNA quality. Sequencing libraries were constructed from 1 µg of total RNA using the NEBNext Ultra II Directional RNA Library Prep Kit for Illumina (NEB). The library construction process included mRNA purification by polyA tail capture with polyT magnetic beads, fragmentation, strand-specific cDNA synthesis, end repair, 3’ end adenylation, adapter ligation, and PCR amplification. The resulting libraries were validated and quantified using Qubit and TapeStation assays. Library preps were pooled in equal ng amounts, and the nM concentration of the pool was verified with an Illumina KAPA Library Quantification qPCR assay 2 (Roche). Each library was indexed with a barcode sequence and sequenced in a multiplexed manner. An Illumina NextSeq 550 system was used to generate single-end, 75-base sequence reads from the libraries. Base calling was performed by the instrument’s Real-Time Analysis (RTA) software. The base call files (bcl files) were demultiplexed and converted to compressed FASTQ files by bcl2fastq2.

### 2.9 RT-qPCR

Real-time quantitative polymerase chain reaction (RT-qPCR) was used to determine the mRNA expression levels of select genes (Table 2) using a Bio-Rad CFX384 real-time PCR detection system and SYBR Green mix (Biomake, B21202).

**Table 2:**
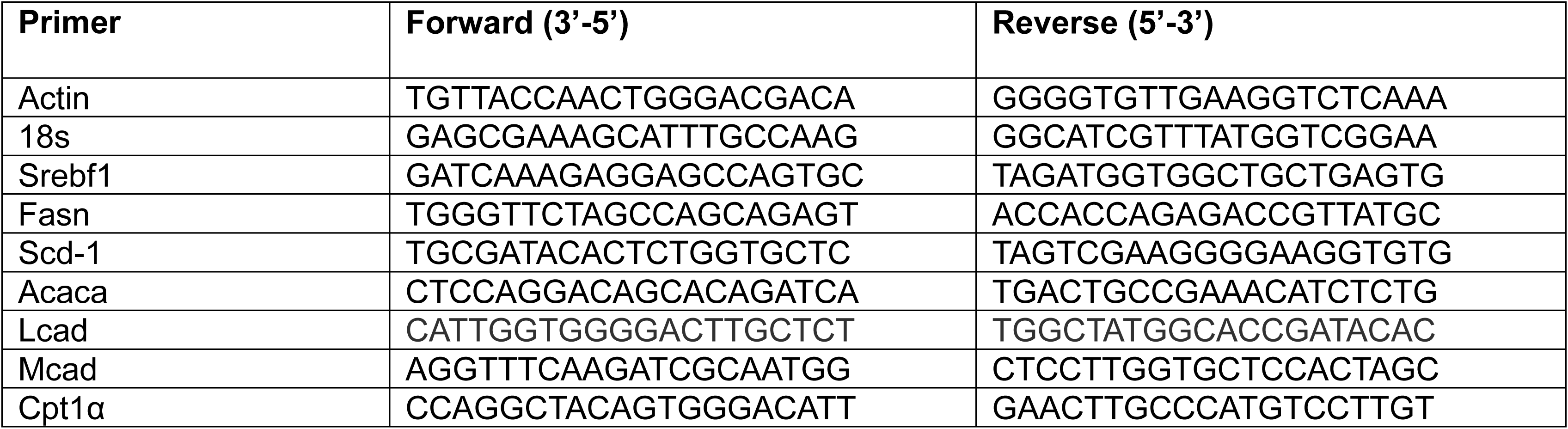
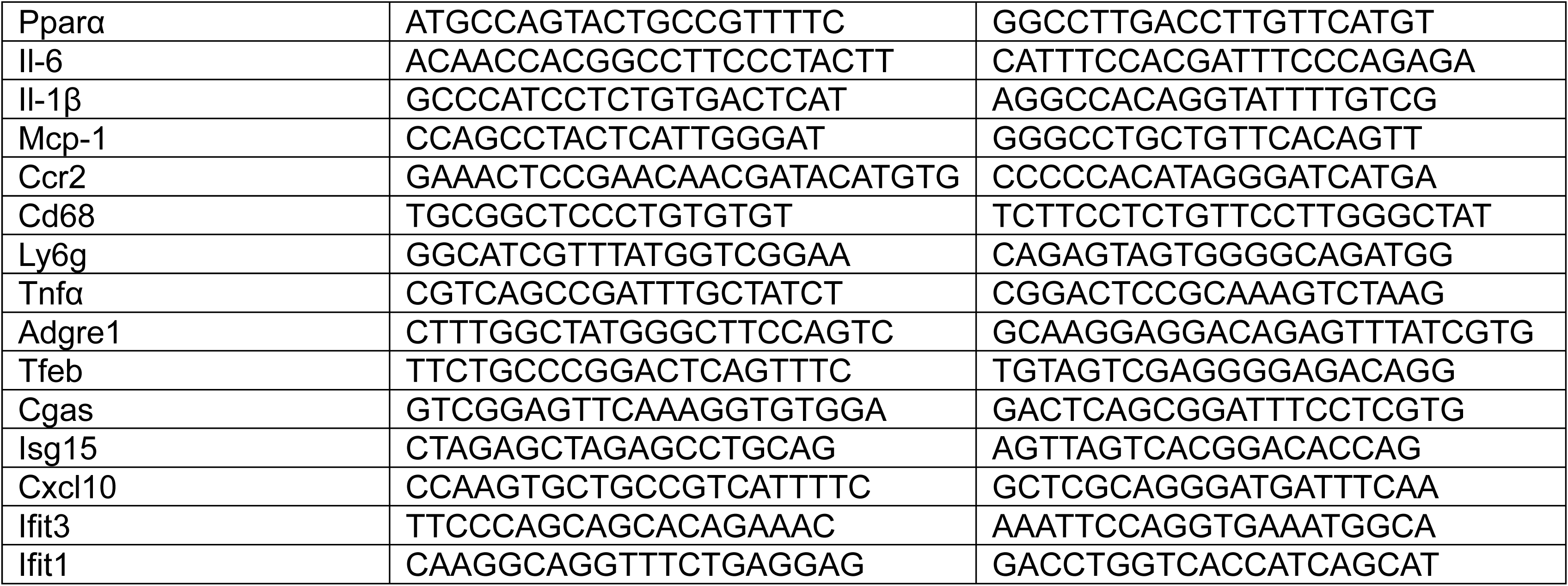
List of Primers (Forward and Reverse) used for RT-qPCR.

### 2.10 Analysis of RNA sequencing results

The RNAseq data were analyzed using our previously established technical pipeline 1, 2. Briefly, we used HISAT2 v.2.1.0.13 to map high-quality reads to the mouse reference genome (GRCm38.90). Gene expression quantification was performed with HTSeq-counts v0.6.0. Significant differentially expressed genes (DEGs) were identified with the R package DESeq2. Statistical significance was assessed using Benjamini-Hochberg-adjusted P values at 0.05. The heatmaps were plotted in R Studio. Principal component analysis (PCA) and volcano plots were prepared in R. Pathway analyses were performed using Ingenuity Pathway Analysis (IPA). We analyzed enrichment of up- and down-regulated genes in the aged control diet-fed group compared with the young control diet-fed group, and in each ethanol diet-fed group compared with the control diet-fed group. Genes with a log2 fold change >2 or <-2 and an FDR-corrected q value <0.05 were used for the analysis.

### 2.11 Metabolomics analysis

Frozen hepatic tissue was submitted to Metabolon for metabolomics analysis. The dataset includes a total of 1031 named and unnamed biochemicals. Statistical analysis was performed using Welch’s two-sample t-test and a p-value of ≤ 0.05 was used for significance.

### 2.12 Proteomics analysis

Serum samples were sent to the University of Arkansas Medical Center (UAMS) for proteomics analysis. Serum samples were analyzed using liquid chromatography/mass spectrometry (LC/MS) on an Orbitrap Exploris™ with data-independent acquisition.

### 2.13 Statistical analysis

Statistical analysis was performed using GraphPad Prism software (version 9.4.1 (681), GraphPad Software, Boston, MA). All experimental data were expressed as mean ± SEM. Analyses were performed using either one-way ANOVA with Tukey’s post hoc test or a two-tailed Student’s t-test, as appropriate. A p-value ≤ 0.05 was considered significant.

## 3. Results

### 3.1 Advanced age drives ethanol-induced triglyceride accumulation, but not liver injury in mice

To assess differences in alcohol-induced liver injury between aged and young mice, serum levels of alanine aminotransferase (ALT) and aspartate aminotransferase (AST) were measured. No differences were observed in serum ALT levels between young and aged mice fed either control or alcohol diets (Figure 1A). Additionally, neither alcohol nor advanced age induced further increases in serum AST levels (Figure 1A). Gao-binge alcohol, combined with advanced age, led to a threefold increase in hepatic triglyceride content (Figure 1C). Although liver injury was not readily evident histologically, lipid accumulation was further confirmed by the presence of large lipid droplets in H&E and Oil-red-O-stained hepatic tissue from aged, alcohol-fed mice compared with young alcohol-fed mice (Figure 1E). To further evaluate alterations in hepatic alcohol metabolism, protein levels of the three primary alcohol-metabolizing enzymes were determined. As expected, Gao-binge alcohol induced an approximately 50% increase in expression of cytochrome P450 2E1 (CYP2E1) in both young and aged livers. Alcohol dehydrogenase (ADH) protein levels remained largely unchanged across the four groups. In young mice fed an ethanol-rich diet, acetaldehyde dehydrogenase-2 (ALDH2) protein levels were two-fold higher than in control diet-fed mice. Although ALDH2 was induced by ethanol in aged, Gao-binge-fed mice, the increase was less pronounced than in young, ethanol-fed mice (Figure 1F-G). Because serum ALT did not show a significant increase, we analyzed ALT protein expression to determine whether age affected ALT protein content in the mouse liver. Western blot analysis revealed that neither age nor increased ethanol consumption substantially affected this protein. Taken together, these data suggest that aged livers are more susceptible to hepatic steatosis but not to liver injury during alcohol consumption.

**Figure 1:**
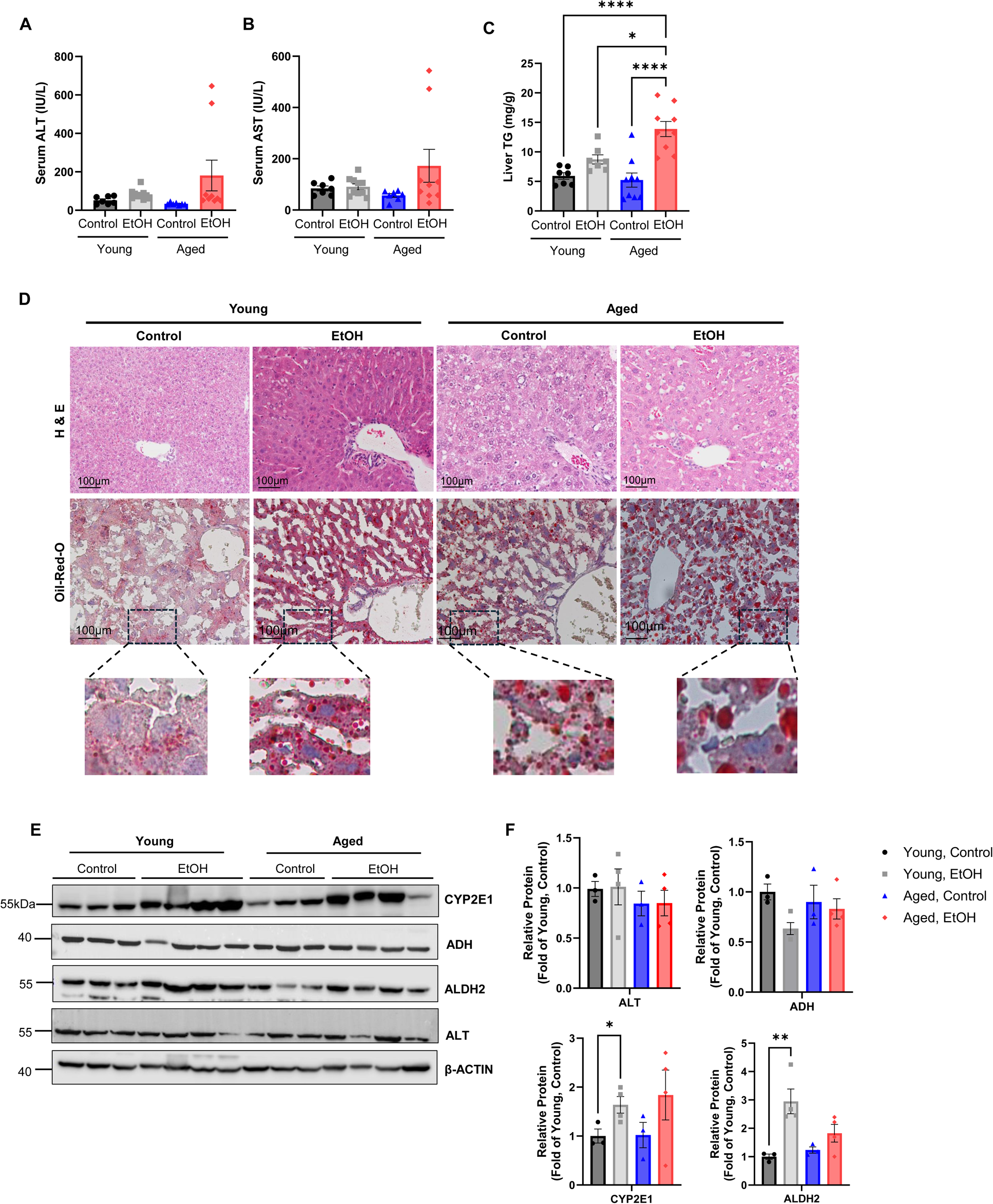
Advanced age drives ethanol-induced triglyceride accumulation, but not liver injury in mice. Young (3-month-old) and aged (22-month-old) C57Bl6/N male mice were subjected to the Gao-binge model. (A-B) Serum ALT and AST levels were measured. (C) Hepatic triglycerides were measured. Data are presented as means± SEM (n=7-9). One-way ANOVA with Tukey’s post hoc test. (D) Representative images of H&E staining and Oil-Red-O staining of liver tissues from control diet and alcohol diet-fed young and aged mice are shown. Lower panel images were enlarged to show lipid droplet size in hepatic tissue. (E) Total liver lysates from each group were used for western blot analysis for the following metabolizing enzymes: cytochrome P450 2E1 (CYP2E1), alcohol dehydrogenase (ADH), acetaldehyde dehydrogenase 2 (ALDH2), alanine transaminase (ALT) (n=3-4). (F) Densitometry analysis of (E) (n=4). Data are presented as means± SEM. Ordinary one-way ANOVA with Tukey’s post hoc test. EtOH, ethanol diet.

### 3.2 Advanced age and Gao-binge ethanol feeding dysregulate the expression of lipid movement and transport genes

To better understand the increased hepatic triglyceride accumulation observed in the livers of aged, alcohol-fed mice, we analyzed RNA sequencing and metabolomic datasets to identify the potential factors contributing to hepatic lipid accumulation. Principal component analysis (PCA) of the RNA-seq dataset showed a clear, distinct separation among all four experimental groups. This observed separation shows that both old age and chronic-on-binge ethanol feeding had distinct effects on the gene expression profile in the liver (Figure 2A). While several genes involved in lipid transport and metabolism had similar basal expression between both young and aged control diet-fed mice, the expression of lipid transport genes like microsomal triglyceride transfer protein (*Mttp*), secretion-associated Ras-related GTPase 1B (*Sar1b*), and apolipoprotein B (*Apob*) had an age-associated decrease in the hepatic tissue of aged, control diet-fed mice (Figure 2B). Advanced age was associated with upregulation of *Cidea* gene expression, which was further increased by ethanol (Figure 2B). Heavy alcohol consumption promoted the upregulation of synthesis genes like diacylglycerol O-acyltransferase 1 (*Dgat1*) and lipin 1 (*Lpin1*), regardless of age, whereas no significant change was observed in the mRNA expression of other genes involved in de novo lipogenesis (DNL) (Figure 2B-C). Additionally, genes involved in lipid transport, such as low-density lipoprotein receptor-related protein 1 (*Lrp1*) and low-density lipoprotein receptor (*Ldlr*), were downregulated by ethanol feeding (Figure 2B).

**Figure 2:**
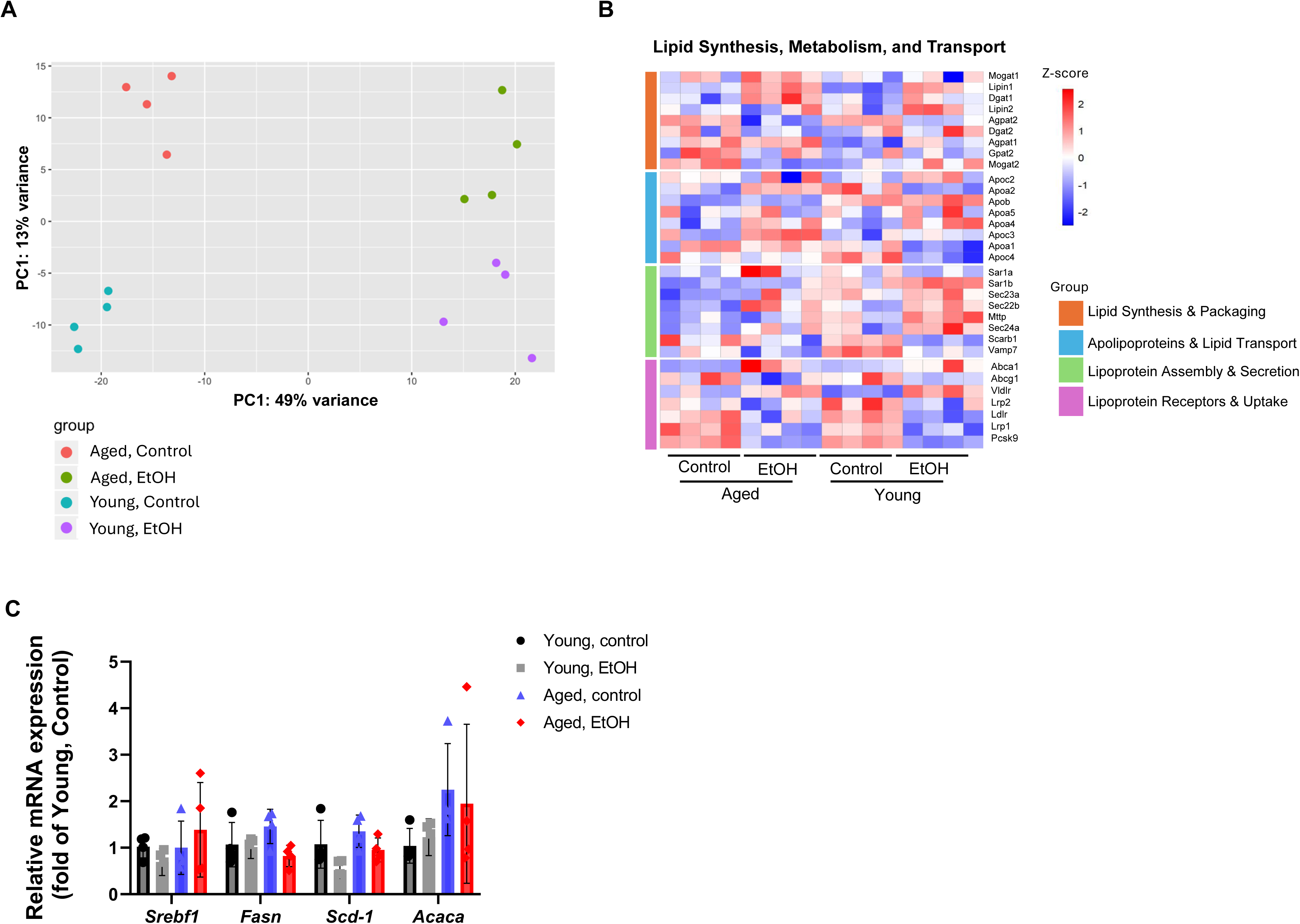
Advanced age and Gao-binge ethanol feeding on the expression of lipid movement and transport genes. Young (3-month-old) and aged (22-month-old) C57Bl6/N male mice were subjected to the Gao-binge model. (A) Principal component analysis (PCA) of RNA sequencing data. (B) Heatmap analysis of genes involved in lipid synthesis, metabolism, and transport from RNAseq dataset (n=4). RNA-seq data can be accessed at GSE 283201. (C) Real-time quantitative PCR (rt-qPCR) analysis of the hepatic mRNA expression of the following *de novo* lipogenesis genes: *Srebf*, *Fasn*, *Scd-1*, and *Acaca* (n=4). Data are presented as means± SEM. Ordinary one-way ANOVA with Tukey’s post hoc test. EtOH, ethanol diet.

### 3.3 Gao-binge ethanol feeding results in increased hepatic concentrations of mono- and diacylglycerol species that are more pronounced in aged livers

Similar to the PCA results from the RNAseq analysis, the metabolomics dataset revealed the same features: a distinct impact of both aging and Gao-binge alcohol feeding on the hepatic metabolite profile (Figure 3A). The hepatic concentration of glycerol-3-phosphate (G3P), the backbone for triacylglycerol synthesis, exhibited a 2-fold increase in hepatic tissue from control diet-fed aged mice compared to the G3P concentrations in hepatic tissue from control diet-fed young mice (Figure 3B, 3D). In contrast, ethanol exposure resulted in a 3-fold decrease in G3P levels in aged, ethanol-fed livers compared to aged, control diet-fed livers (Figure 3B).

**Figure 3:**
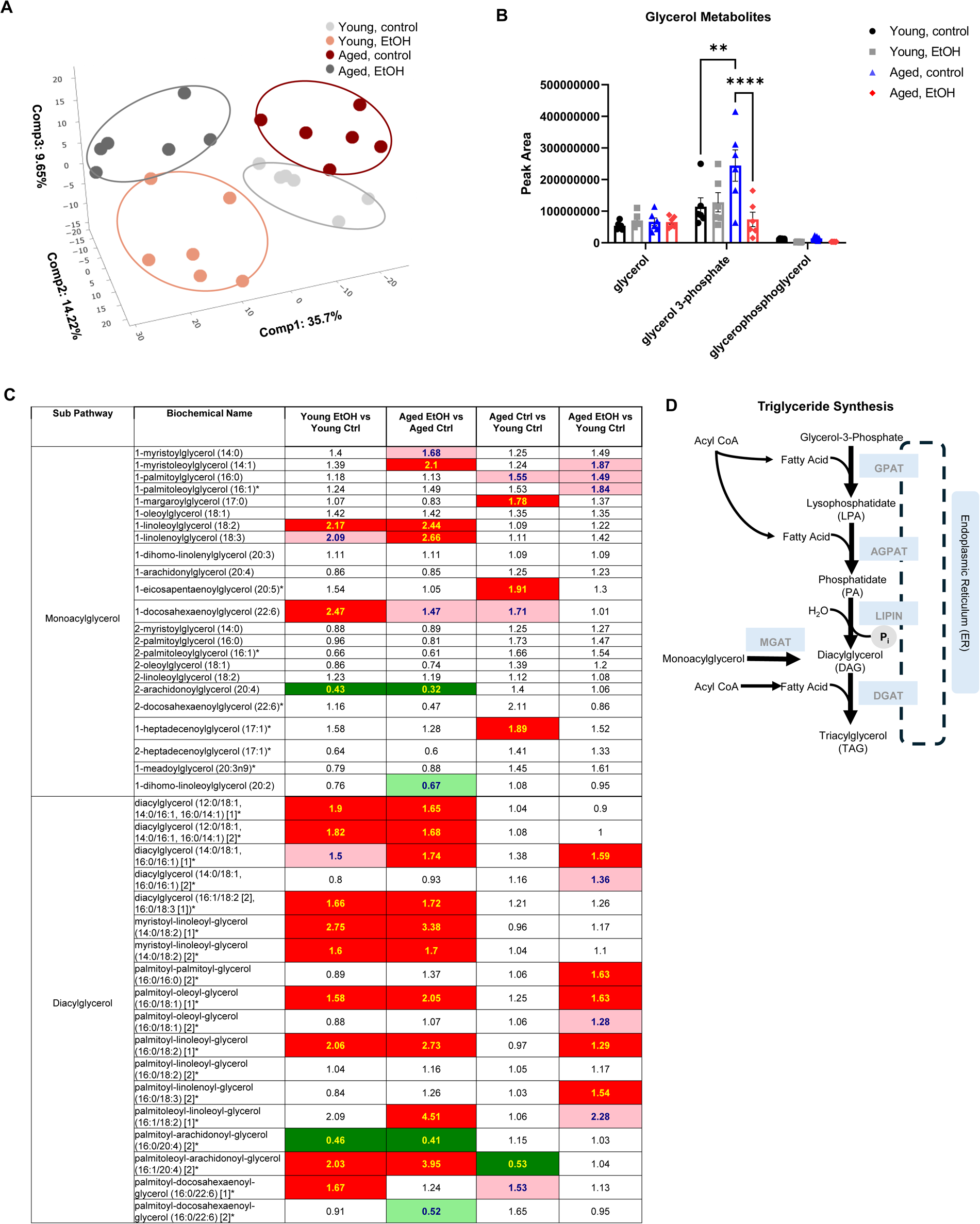
Gao-binge ethanol feeding results in increased hepatic concentrations of mono- and diacylglycerol species that are more pronounced in aged livers. Young (3-month-old) and aged (22-month-old) C57Bl6/N male mice were subjected to the Gao-binge model. (A) PCA of metabolomics dataset (n=6). (B) Quantification of hepatic glycerol metabolites (n=6). Data are presented as means± SEM. Ordinary one-way ANOVA with Tukey’s post hoc test. (C) Heatmap analysis of mono- and diacylglycerol species from metabolomics analysis (n=6). (D) Schematic of the triglyceride synthesis pathway. **, p>0.01; ****, p>0.0001. Ctrl, control diet; EtOH, ethanol diet.

Furthermore, ethanol feeding in young mice resulted in a marked increase in monoacylglycerol (MAGs) and diacylglycerol (DAGs) species, lipid species used in the synthesis of triglycerides. This change was more pronounced in the livers of aged, Gao-binge-fed mice (Figure 3C, 3D). These results indicate that ethanol feeding and aging increased some intermediate triglyceride synthesis metabolites in the mouse livers.

### 3.4 Old age mice do not have impairment in fatty acid beta-oxidation induced by Gao-binge alcohol

Heatmap analysis showed an increased trend in the expression of several genes associated with fatty acid (FA) metabolism due to Gao-binge alcohol feeding, regardless of age (Figure 4A). Genes involved in peroxisomal FA breakdown and transport, such as Eci2, Phyh, and Slc27a2, were downregulated in aged livers, irrespective of alcohol feeding (Figure 4A). No change was observed in the mRNA levels of mitochondrial FA-metabolizing genes. However, mRNA expression of peroxisome proliferator-activated receptor-alpha (Pparα) was decreased to some extent in aged, alcohol-fed mice but did not reach statistical significance (Figure 4B). There was a significant increase in numerous medium-chain and long-chain saturated and monounsaturated acylcarnitine metabolites in aged, alcohol-fed mice compared with young, alcohol-fed mice (Figure 4C). Additionally, metabolomics analysis showed a massive increase in hepatic 3-hydroxybutyrate (BHBA) levels in Gao-binge ethanol-fed young mice, with even higher levels of BHBA in ethanol-fed aged mice (Figure 4D), suggesting that age and ethanol-feeding do not impair fatty acid beta oxidation.

**Figure 4:**
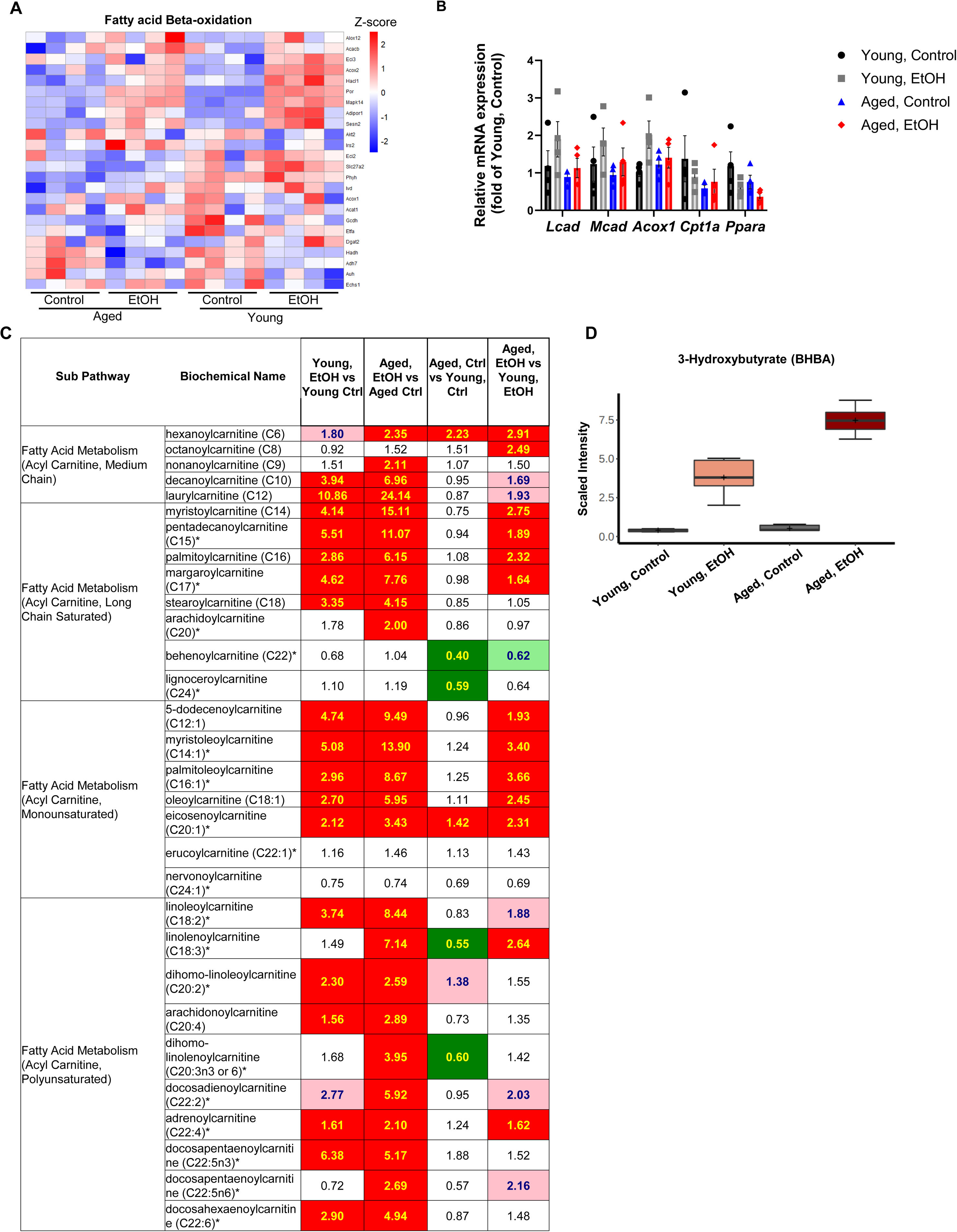
Old age does not impair fatty acid beta oxidation induced by Gao-binge alcohol. Young (3-month-old) and aged (22-month-old) C57Bl6/N male mice were subjected to the Gao-binge model. (A) Heatmap analysis of fatty acid β-oxidation genes from RNAseq dataset for each experimental group (n=4). (B) rt-qPCR results of the hepatic mRNA expression of the following genes involved in fatty acid metabolism: *Lcad*, *Mcad*, *Acox1*, *Cpt1α*, and *Pparα* (n=4). Data are presented as means± SEM. Ordinary one-way ANOVA with Tukey’s post hoc test. (C) Heatmap analysis of hepatic acyl carnitine metabolite species from metabolomics dataset (n=6). (D) Quantification of hepatic content of 3-hydroxybutyrate (BHBA) from metabolomics dataset (n=6).

### 3.5 Advanced age and Gao-binge ethanol feeding synergistically promote hepatic senescence

To detect changes in the regulation of senescence during aging and Gao-binge alcohol feeding, we analyzed our RNAseq dataset and found induction of genes associated with the inhibition of senescence in response to stress, such as *Fos, Rps6ka3*, and *Ubc,* whereas genes involved in cell cycle progression, like *Cdk4* and *Cdk6,* showed decreased expression. Ethanol feeding upregulated the expression of senescence-promoting genes, such as *Cdc27, Igfp7, Cdkn1a,* and *Cdkn1b* (Figure 5A).

**Figure 5:**
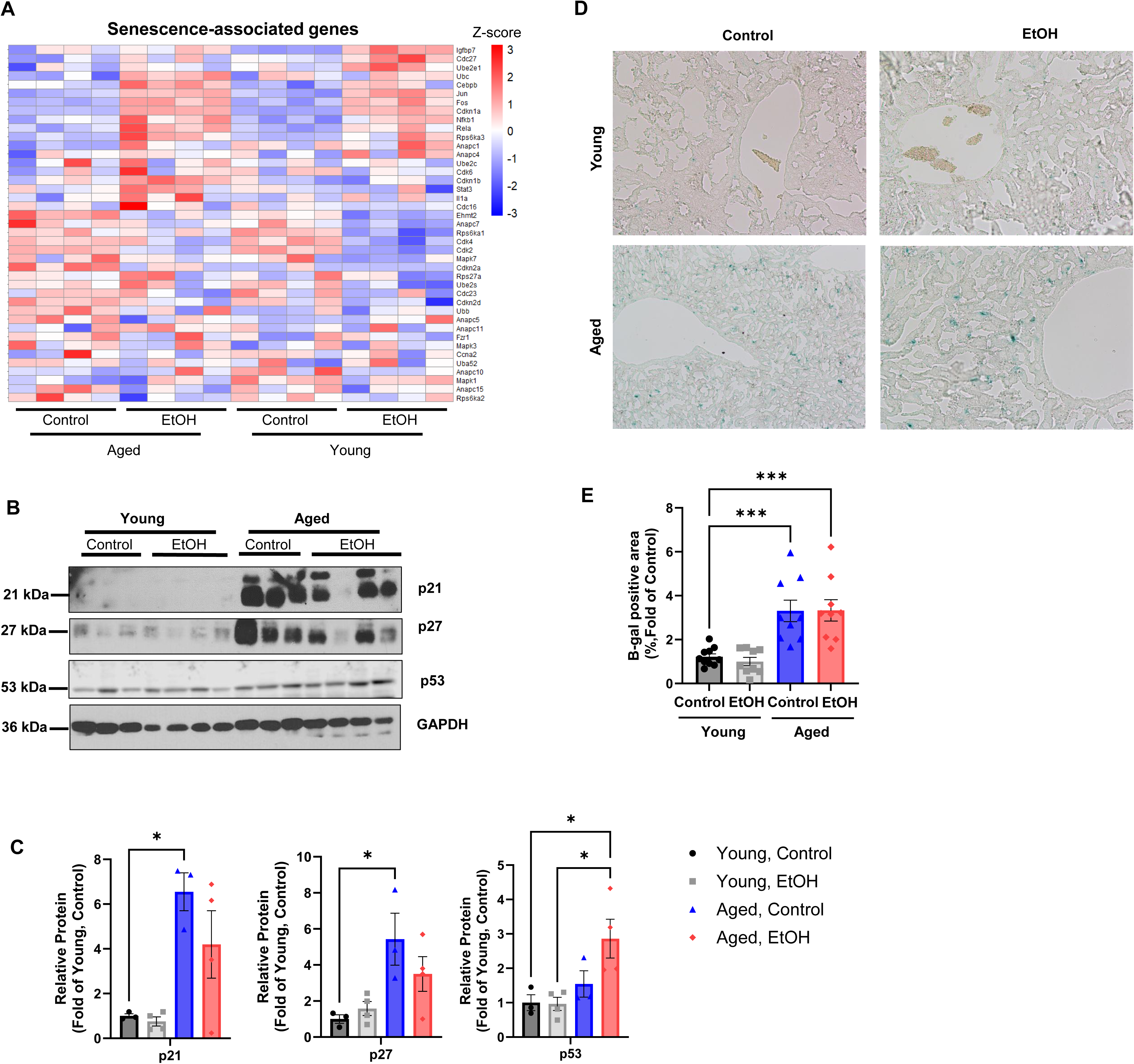
Advanced age and Gao-binge ethanol feeding synergistically promote hepatic senescence. Young (3-month-old) and aged (22-month-old) C57Bl6/N male mice were subjected to the Gao-binge model. Heatmap analysis of senescence-associated genes from RNAseq data for each experimental group (n=4). (B)Total liver lysates were subjected to western blot analysis. Representative blots for the expression of the following proteins: p21, p27, and p53 (n=3-4). (C) Densitometry analysis of (B) (n=3-4). (D) Representative images of senescence-associated β-galactosidase activity staining in liver tissue. (E) Quantification of (D) (n=2). Data are presented as means± SEM. Ordinary one-way ANOVA with Tukey’s post hoc test. *, p>0.02; ***, p>0.0005. EtOH, ethanol diet.

In exploring the observed induction of cellular senescence by Gao-binge feeding and aging, we assessed the protein expression of p21 and p27, which are encoded by *Cdkn1a* and *Cdkn1b,* respectively. Both p27 and p21 protein levels were increased in the livers of aged, ethanol-fed mice, aligning with our RNAseq data (Figure 5B-C). In conjunction with elevated p21 and p27 protein levels, aging and Gao-binge ethanol feeding resulted in a 2-fold increase in p53 protein in comparison to Gao-binge-fed young mice (Figure 5B-C). Additionally, we observed an increased number of positively stained foci for senescence-associated beta-galactosidase (SA β-Gal) in liver tissue from both aged control-diet and ethanol-fed mice (Figure 5D-E). These results indicate that both advanced age and excessive ethanol intake promote cellular senescence in the liver.

### 3.6 Gao-binge ethanol feeding induces pro-inflammatory signaling that is perpetuated by old age

Immunostaining for Kupffer cell (KC) using the marker F4/80 and neutrophil recruitment using the surface marker Ly6G did not reveal any significant differences in the positively stained areas in liver tissue from both control diet and ethanol-fed young mice (Figure 6A-B). On average, aged mice exhibited approximately twice as much F4/80+ staining as young mice, regardless of diet. Aging prompted a significant increase in positive Ly6G foci in hepatic sections from control diet, aged mice, compared to young mice fed a similar diet. The presence of Gao-binge ethanol feeding did not further elevate the numbers of positive foci observed in aged mice (Figure 6A-B). Old age was associated with higher gene expression of chemokines such as C-C chemokine receptor type-2 (*Ccr2*) and chemokine ligand-1 (*Cxcl1*) (Figure 6C). The increased expression of *Ccr2* in ethanol-fed aged mice was further confirmed by qPCR (Figure 6D). Gao-binge alcohol feeding upregulated other inflammatory genes, such as *Cd14* and *Ccl6*, in both young and aged mice compared with their age-matched control diet-fed groups (Figure 6C).

**Figure 6:**
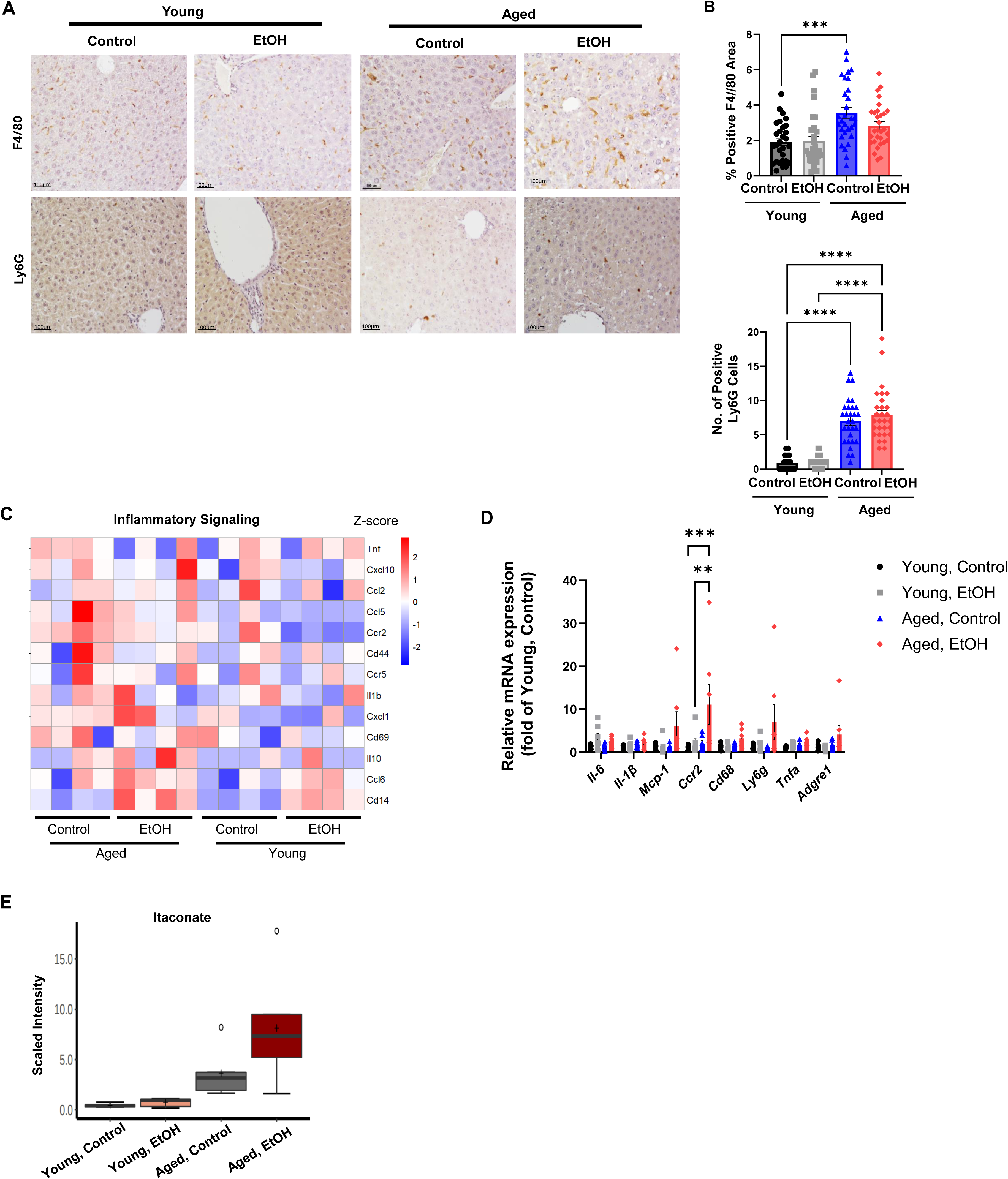
Gao-binge ethanol feeding induces pro-inflammatory signaling that is perpetuated by old age. Young (3-month-old) and aged (22-month-old) C57Bl6/N male mice were subjected to the Gao-binge model. (A) Representative images of F4/80 and Ly6G staining in hepatic tissue from each experimental group (n=3). (B) Quantification of (A) (n=3). (C) Heatmap analysis of select pro-inflammatory genes, cytokines, and chemokines for each experimental group (n=4). (D) rt-qPCR results of the hepatic mRNA expression of the following pro-inflammatory genes, cytokines, and chemokines: *Il-6*, *Il-1β*, *Mcp-1*, *Ccr2*, *Cd68*, *Ly6g*, *Tnfα, Adgre1* (n=6-7). (E) Quantification of hepatic itaconate content from metabolomics dataset (n=6). Data are presented as means± SEM (B, D). Ordinary one-way ANOVA with Tukey’s post hoc test (B, D). **, p>0.003; ***, p>0.0003; ****, p>0.0001. EtOH, ethanol diet.

Furthermore, untargeted metabolomic analysis of hepatic tissue from the four experimental animal groups revealed that advanced age and Gao-binge alcohol synergistically drove a marked increase in hepatic itaconate (Figure 6E), a metabolite that indicates increased inflammation. Together, these data indicate that old age further exacerbates alcohol-induced inflammatory signaling in the liver.

### 3.7 Old age and Gao-binge ethanol feeding synergistically promote upregulation of innate immune pathways of IL-7, IFNs, and cGAS while prompting increased secretion of inflammatory molecules

Pathway analysis of our genomic dataset revealed that advanced age as well as the combination of old age and Gao-binge ethanol feeding upregulated interleukin-7 (IL-7) and interferon signaling pathways (Figure 7A). To further investigate the upregulation of these two pathways, heatmap analysis of interferon and interleukin genes as well as regulatory genes involved in the cGAS (cyclic GMP-AMP synthase)-STING (stimulator of interferon genes) pathway showed that chronic-on-binge ethanol feeding paired with advanced age drove an increased in the expression of genes like *interferon-3* (*Irf3*) and *interferon-7* (*Irf7*) as well as IFN-stimulated genes such as *interferon-induced proteins with tetratricopeptide repeats* (*Ifit1*, *Ifit2*, *Ifit3*) (Figure 7B). Untargeted proteomic analysis of serum samples from the four experimental animal groups showed that advanced age and Gao-binge alcohol synergistically prompted the secretion of pro-inflammatory proteins such as immunoglobin protein fragment variants of IGKV4 and anti-inflammatory enzymes like leukotriene A4 hydrolase (LTA4H) and serine protease inhibitor (SERPINA3J) in aged, alcohol-fed mice (Figure 7C). Additionally, analysis of the serum concentrations of various inflammatory molecules using a cytokine/chemokine array displayed higher levels of chemokine C-C motif ligand 6 (CCL6) in the serum from young, ethanol-fed mice, aged, control diet fed, and aged, ethanol-fed mice (Figure 7D-E). Other inflammatory-associated molecules such as insulin-like growth factor-binding protein-1 (IGFBP-1) and low-density lipoprotein-receptor (LDL-R) were observed to be higher due to Gao-binge alcohol and aging as well (Figure 7D-E). These results highlight the synergistic impact of old age and ethanol feeding on elevating inflammatory pathways in the liver.

**Figure 7:**
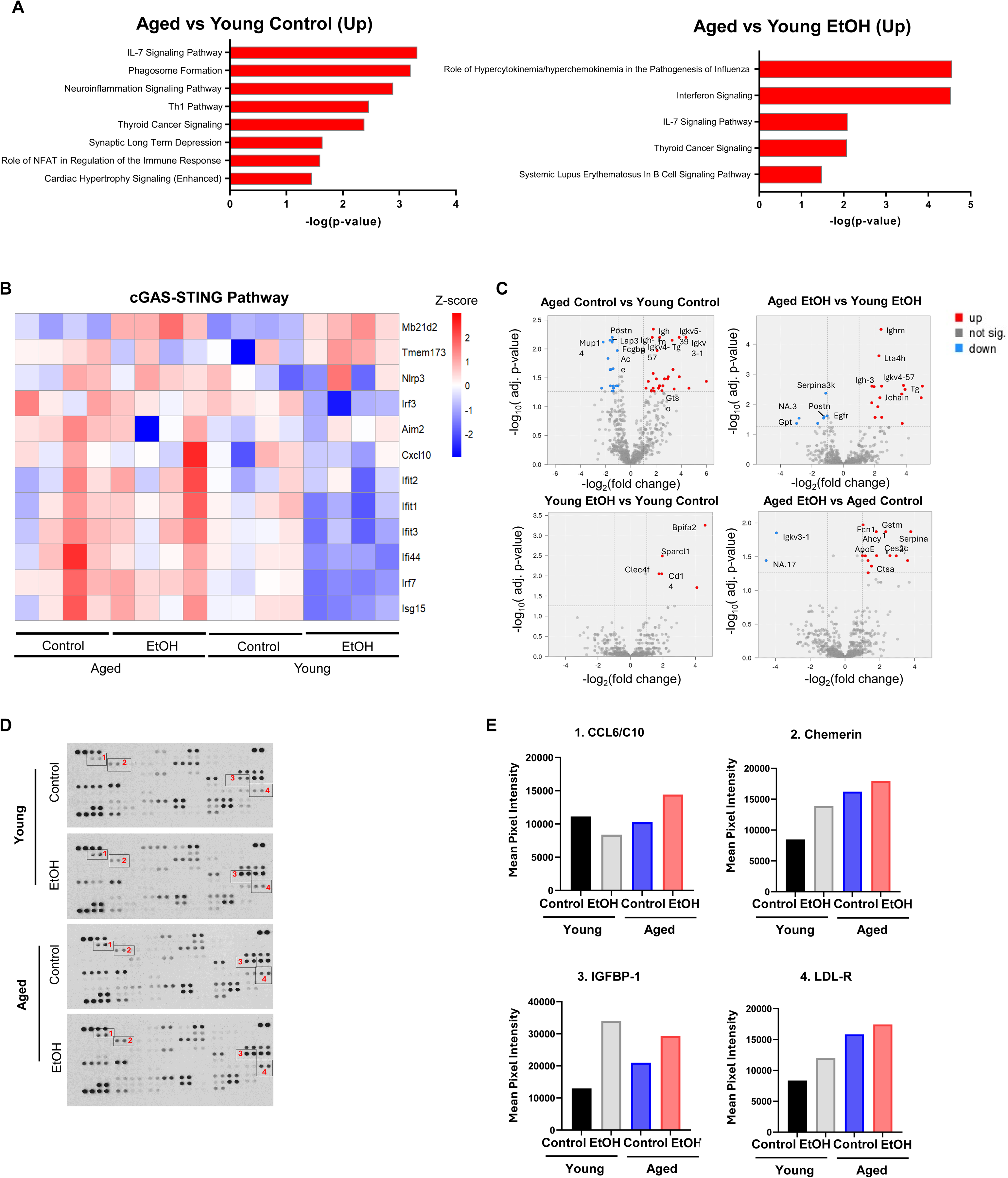
Old age and Gao-binge ethanol feeding synergistically promote upregulation of innate immune pathways of IL-7, IFNs, and cGAS while prompting increased secretion of inflammatory molecules. Young (3-month-old) and aged (22-month-old) C57Bl6/N male mice were subjected to the Gao-binge model. (A) Pathway analysis from the RNAseq dataset for stated group comparison. (B) Heatmap analysis of genes in cGAS-STING pathway from the RNAseq dataset (n=4). (C) Volcano plots for selected comparisons from serum proteomics analysis. (D) Representative blots of cytokine/chemokine array for each experimental group. Black boxes with numbers highlight changes in density of selected cytokines. (E) Quantification of the density results from the selected cytokines highlighted by black boxes.

### 3.8 Advanced age and Gao-binge ethanol feeding result in reduced TFEB-mediated autophagy

We have previously shown that autophagy mediated by transcription factor-EB (TFEB) is impaired by Gao-binge alcohol feeding in mice (Chao, Wang et al. 2018). Western blot and qPCR analysis showed that hepatic TFEB levels were similar in both young and aged control diet-fed mice whereas alcohol feeding led to a moderate reduction of hepatic TFEB (Figure 8A-B, 8F). Gao-binge ethanol feeding markedly increased the levels of microtubule-associated 1 light chain 3-II (LC3-II) with a moderate decrease of p62 in young mice. Notably Gao-binge ethanol feeding did not alter the levels of LC3-II compared with the control-diet fed mice.

**Figure 8:**
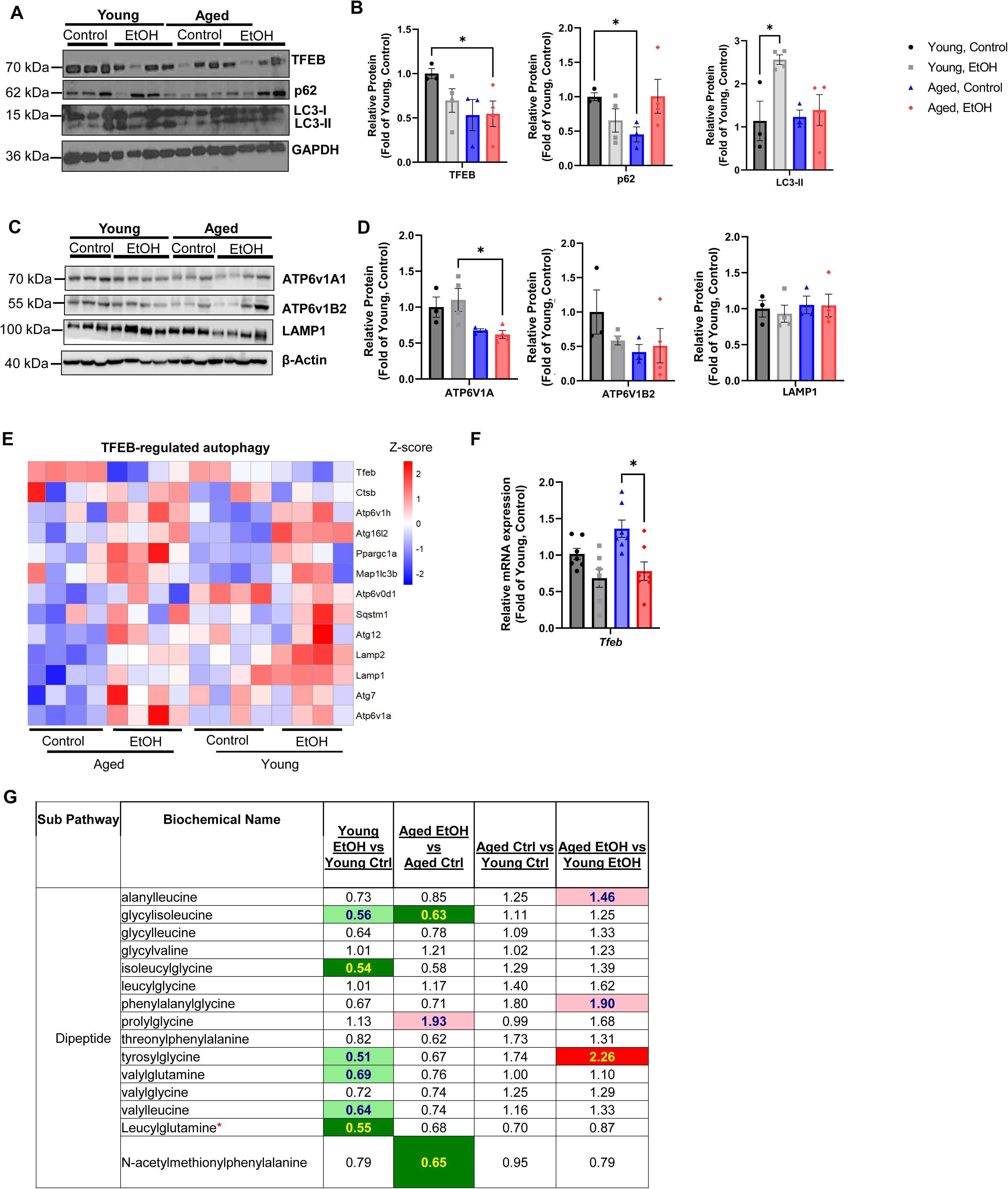
Advanced age and Gao-binge ethanol on hepatic TFEB and autophagy. Young (3-month-old) and aged (22-month-old) C57Bl6/N male mice were subjected to the Gao-binge model. (A) Total liver lysates were used for western blot analysis for the following autophagy proteins: TFEB, p62, and LC3 (I-II) (n=3-4). (B) Quantification of (A) (n=3-4). (C) Representative immunoblot of the lysosomal proteins: ATP6v1A1, ATP6v1B2, and LAMP1 (n=3-4). (D) Quantification of (C) (n=3-4). (E) Heatmap analysis of TFEB-regulated autophagic genes from the RNAseq dataset for each experimental group (n=4). (F) rt-qPCR results of the hepatic mRNA expression of *Tfeb* (n=7). (G) Heatmap analysis of hepatic dipeptide metabolites from metabolomics dataset (n=6). Data are presented as means± SEM (B, D, F). Ordinary one-way ANOVA with Tukey’s post hoc test (B, D, F). *, p>0.05. EtOH, ethanol diet.

The basal levels of p62 were already lower in aged mice compared with young mice but ethanol feeding further increased p62 levels in aged mice (Fig 8A-B). Aged livers showed ∼25% and 50% lower protein expression of the lysosomal ATPases ATPase H+ transporting V1 subunit A1 (ATP6V1A1) and ATPase H+ transporting V1 subunit B2 (ATP6V1B2), respectively, whereas lysosome-associated membrane protein-1 (LAMP1) content remained largely unchanged across the four experimental groups (Figure 8C-D). RNAseq analysis showed that ethanol feeding decreased *Tfeb* mRNA levels in both young and aged mice (Figure 8E-F). However, overall TFEB signature/target gene expression was unchanged in ethanol-fed young mice but increased in aged mice (Figure 8E). Notably, advanced age was associated with lower basal expression of Atp6v1a and Atp6v0d1 compared with young mice (Figure 8E). Dipeptides are generated from the lysosomal degradation of peptides and may be used as a marker for lysosomal degradation activity. Analysis of the metabolomics dataset showed that ethanol feeding decreased dipeptide production in both young and aged livers, with more pronounced defects in young mice (Figure 8G). Taken together, these results suggest that ethanol may negatively affect lysosomal degradation, and that aging appears to have little synergistic effect in further impairing lysosomal functions compared with ethanol feeding in mouse livers.

### 3.9 Hepatic overexpression of TFEB reduced immune cell activation and infiltration in aged, Gao-binge ethanol-fed mice

We next determined whether overexpression of TFEB in aged mouse livers would protect against Gao-binge ethanol-induced liver injury. Overexpression of TFEB decreased serum ALT and AST levels to approximately 50% and 20%, respectively, in ethanol-fed aged mice compared with mice that received only a control virus, although the changes did not reach statistical significance. However, a correlational analysis showed that higher hepatic TFEB mRNA levels were associated with lower serum ALT (Figure 9A-C). No significant differences in hepatic TG levels or histology were observed between ethanol-fed control mice (AD-Null) and those with TFEB overexpression (Figure 9D-E). In contrast, TFEB overexpression markedly reduced F4/80+ and Ly6G+ staining in aged, Gao-binge ethanol-fed livers compared with the AD-Null group (Figure 9F-G). Furthermore, TFEB overexpression resulted in a ∼50-70% reduction in IRF-7, cGAS, and IRF-3 protein levels and significantly decreased caspase-1 activity compared with control mice (Figure 9H-J). Similar to the protein changes, the mRNA levels of cGAS-STNG-related genes, such as *cGas, Isg15, Cxcl10, Ifit3,* and *Ifit1*, also tended to be lower in mice with TFEB overexpression compared with control mice (Figure 9K). Taken together, these data suggest that increased hepatic TFEB content may play a role in reducing age- and alcohol-induced inflammation but not steatosis.

**Figure 9:**
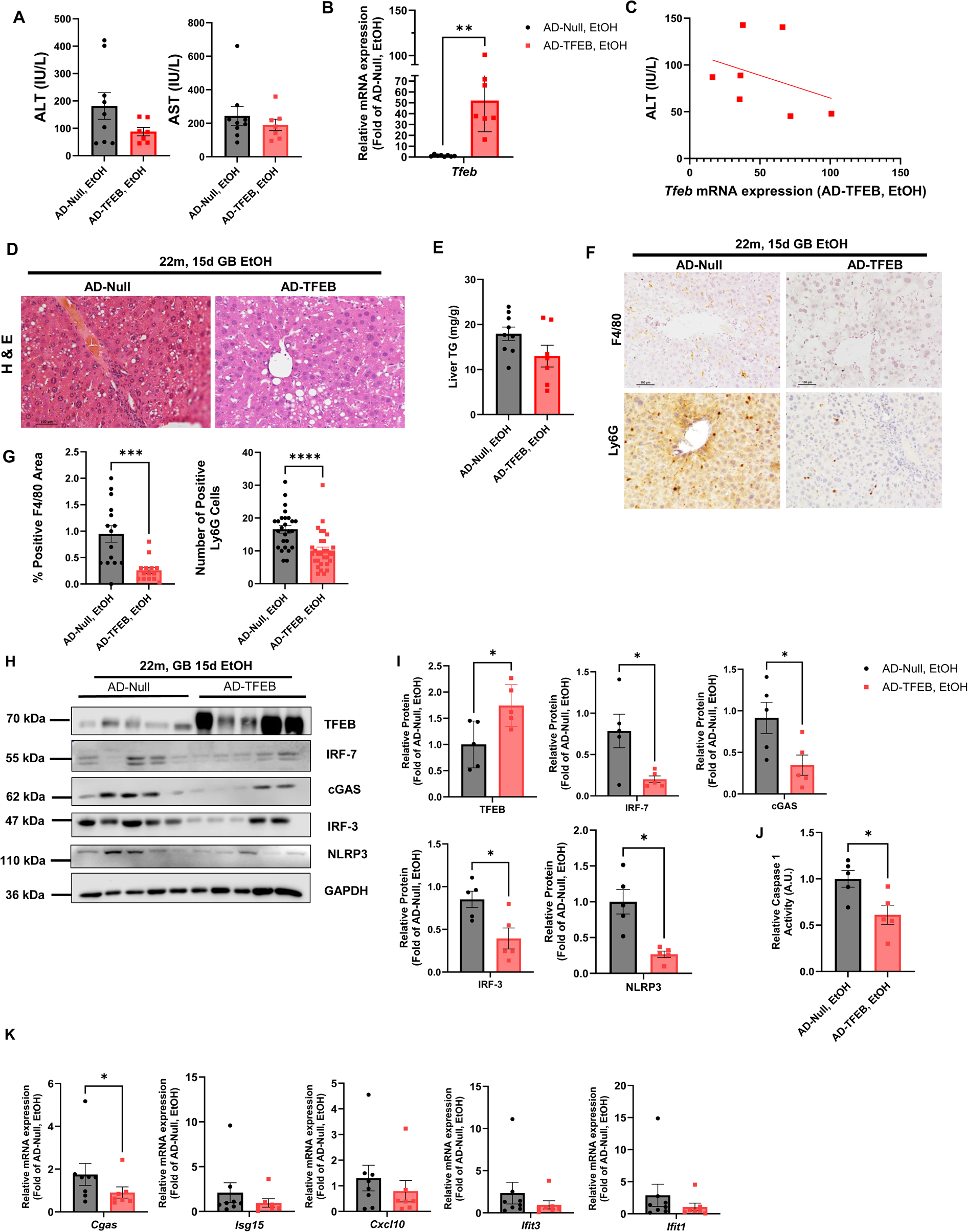
Hepatic overexpression of TFEB reduced cGAS-STING-mediated innate immune response and immune cell infiltration in Gao-binge ethanol-fed aged mice. Aged (22-month-old) C57Bl6/N male mice received tail-vein injections with adenovirus (AD) to overexpress TFEB in the liver then were subjected to the Gao-binge model. (A) Serum ALT and AST levels were measured (n=7-9). (B) rt-qPCR results of hepatic *Tfeb* mRNA expression (n=7-9). (C) Correlation graph of ALT to *Tfeb* mRNA expression (n=7). (D) Representative H&E images of hepatic tissue for AD-Null and AD-TFEB aged mice. (E) Measurement of hepatic triglycerides (n=7-9). (F) Representative immunostaining images for F4/80 and Ly6G (n=3). (G) Quantification of (F). (H) Representative immunoblots of the following proteins: TFEB, IRF-7, cGAS, IRF-3, and NLRP3 (n=5). (I) Quantification of (H) (n=5). (J) Quantification of relative caspase 1 activity (n=5). (K) rt-qPCR results of the hepatic mRNA expression of the following genes: *Cgas*, *Isg15*, *Cxcl10*, *Ifit3*, and *Ifit1* (n=7-8). Data are presented as means± SEM (A-B, E, G, I-K). Ordinary one-way ANOVA with Tukey’s post hoc test (A-B, E, G, I-K). *, p>0.05; **, p>0.003; ***, p>0.0007, ****, p>0.0001. EtOH, ethanol diet.

## Discussion

Old age, or aging, contributes to the development of chronic diseases due to the gradual decline in biological functions essential to homeostasis and overall stress response. The detrimental factors that influence aging have been collectively identified as the “hallmarks of aging” and include 1) genomic instability, 2) telomere attrition, 3) epigenetic alterations, 4) loss of proteostasis, 5) impaired macro-autophagy, 6) dysbiosis, 7) chronic inflammation, 8) disrupted cellular communication, 9) stem cell exhaustion, 10) dysregulated nutrient sensing, 11) mitochondrial dysfunction, and 12) cellular senescence (Lopez-Otin, Blasco et al. 2023). In the liver, aging is associated with a 20-40% decline in liver volume, which can negatively impact overall hepatic metabolic function. Additionally, decreased regenerative capacity of the liver as well as the persistent production of oxidative stress, have been noted to occur with aging (Timchenko 2009, Wang 2021, Williams and Ding 2025). In the present study, using a clinically relevant Gao-binge alcohol feeding model (Cao, Chao et al. 2026), we showed that the combination of aging and alcohol mainly potentiated hepatic steatosis and inflammation but had a moderating role in liver injury.

The liver is a dynamic organ that performs several functions, including regulating lipid metabolism by facilitating fatty acid synthesis and oxidation, as well as exporting fatty acids from the liver for utilization by other tissues (Nguyen, Leray et al. 2008). Excessive alcohol intake impairs these processes, leading to increased fat deposits, or lipid droplets, in the liver (Jeon and Carr 2020, Mandrekar and Mandal 2024). Moreover, proteins associated with DNL and FA metabolism, such as CD36 (cluster of differentiation 36) and CPT1α (carnitine palmitoyltransferase 1A), as well as the transcriptional regulator PPARα are noted to be altered. One of the most remarkable changes in our study was increased hepatic steatosis in ethanol-fed aged mice compared with young mice. RNA-seq analysis did not show significant changes in gene expression in DNL, lipid transport, lipoprotein secretion, and fatty acid beta-oxidation. There was an increase in both medium-chain and long-chain acyl carnitines and in hepatic BHBA levels, likely due to increased fatty acid beta-oxidation as an adaptive response. Interestingly, hepatic BHBA levels were even higher in ethanol-fed aged mice, suggesting that aging does not impair fatty acid beta-oxidation. We previously showed that ethanol feeding increased the number of hepatic megamitochondria and fatty acid beta-oxidation as an adaptive response (Ma, Chen et al. 2023, Niu, Wei et al. 2026). Future studies are needed to investigate whether aging would affect hepatic megamitochondria. However, ethanol administration increased the lipid substrates glycerol-3-phosphate and diacylglycerols in ethanol-fed aged mice, which may partially contribute to increased triglyceride synthesis in these mice. Future studies should use stable isotope tracers and monitor isotopic enrichment in TGs over time to calculate synthesis rates, turnover, and flux through pathways such as de novo lipogenesis or fatty acid uptake.

Perhaps another significant finding in the present study was an increased inflammatory response in ethanol-fed aged mice compared with young mice. In the liver, reactive oxygen species (ROS) produced from ethanol metabolism and endotoxin release from the intestine will activate Kupffer cells (KCs), stimulating nuclear factor-kappa beta (NF-κβ) and the downstream production and release of inflammatory signaling molecules such as tumor necrosis factor-alpha (TNFα), interleukin-8 (IL-8), and monocyte chemoattractant protein-1 (MCP-1). Additionally, cytokine release stimulates the recruitment of circulating neutrophils and macrophages to the liver, further perpetuating hepatocyte damage (Xu and Wang 2022, Dukic, Radonjic et al. 2023, Hong, Huang et al. 2024). Chronic inflammation, or inflammaging, in cells results from the persistent presence of hallmarks of aging, which trigger the continuous production of pro-inflammatory cytokines and ROS. This reaction ultimately induces cellular senescence, or permanent cell cycle arrest (Uyar, Palmer et al. 2020, Baechle, Chen et al. 2023, Lopez-Otin, Blasco et al. 2023). Furthermore, senescent cells will secrete factors associated with senescence, collectively termed the senescence-associated secretory phenotype (SASP), which will prime and promote cell cycle arrest in neighboring cells (Uyar, Palmer et al. 2020, Baechle, Chen et al. 2023, Ajoolabady, Pratico et al. 2025). It has been reported that hepatocyte senescence is attributable to excessive alcohol consumption. Liver sections from patients at various stages of ALD showed increased p21 expression, a cell cycle inhibitor and a marker of senescence (Aravinthan, Pietrosi et al. 2013). This observation was recapitulated in animal studies using chronic ethanol administration models, in which increased p16 and senescence-associated beta-galactosidase (SA β-Gal) were present, along with elevated levels of p21 (Wan, McDaniel et al. 2017, Tian, Xue et al. 2025). In the present study, we also found increased senescence markers (p21 and p27) and SA β-Gal staining in ethanol-fed aged mouse livers, as well as increased secretion of both inflammatory and senescence-associated proteins in the serum. This may synergistically contribute to increased pro-inflammatory signaling within hepatic tissue and to heightened macrophage infiltration in the aged liver, driven by ethanol and aging. Additionally, we found increased cGAS-STING and inflammasome-mediated innate immune responses in eth’[=,manol-fed aged mice. Together, aging, ethanol-induced hepatocyte senescence, and cGAS-STING-mediated innate immune response may contribute to increased hepatic inflammation. Future studies are needed to determine the mechanisms of CGAS-STING activation and whether targeting the cGAS-STING pathway would improve inflammation in aging-associated ALD.

The evolutionarily conserved process of autophagy has long been studied in aging research. It has been thought that autophagy declines with aging and that pharmacological or genetic increases in autophagy may extend lifespan in *C. elegans* and mice (Pyo, Yoo et al. 2013, Kitada and Koya 2021). TFEB, the master regulator of lysosomal biogenesis, is also a central regulator of aging, as its activation extends lifespan and healthspan in flies, nematodes, and mice (Abokyi, Ghartey-Kwansah et al. 2023). In the liver, it has been suggested that a decline in autophagy is attributable to an age-related reduction in LC3 protein and autophagosomal vesicles (Uddin, Nishio et al. 2012, Williams and Ding 2025). However, whether alcohol impairs hepatic autophagy and TFEB in ALD is complex due to several reasons.

First, autophagy is highly dynamic, and there are few reliable, quantitative assays that accurately capture its temporal and spatial dynamics in the liver after alcohol consumption. Although adding a lysosomal inhibitor can help assess autophagic flux in mouse livers, it generally reflects only a single time point because it is impractical to obtain multiple time points in mouse livers after alcohol consumption. Therefore, conclusions about autophagy activity after alcohol consumption should be drawn with caution, as most studies rely solely on steady-state changes in autophagy-related proteins or on a single time point of autophagic flux. Currently, there are also no data showing spatial zonal changes in autophagic activity in mouse livers after alcohol consumption. Second, while RNA-seq analysis can provide extensive information on gene expression changes, including those in autophagy-related genes, after alcohol consumption, mRNA changes often do not correlate with protein expression. In our present study, we did not observe significant, robust changes in hepatic mRNA levels of autophagy-related genes and TFEB signature genes between alcohol-fed old and young mice. Levels of TFEB mRNA and protein also did not differ between alcohol-fed young and aged mice, although the protein level of ATP6V1A was lower in ethanol-fed aged mice. Nevertheless, decreased hepatic dipeptide levels in ethanol-fed mice may suggest reduced lysosomal degradation capacity. Because there was no difference in dipeptide levels between young and aged mice, this suggests that alcohol, rather than aging, may be more critical for impairing lysosomal degradation. However, as discussed above, owing to the complexity and limitations of autophagic flux assays in vivo, more studies with lysosomal inhibitors and multiple time points are needed to more robustly assess autophagy and lysosomal activities in ethanol-fed aged mice. Despite limitations in accurately assessing autophagic activity in mouse livers, pharmacological and genetic activation of TFEB protects against alcohol-induced liver injury in the Gao-binge model (Chao, Wang et al. 2018). We also found that moderate TFEB overexpression protects against alcohol-induced liver injury and markedly improves the cGAS-STING-mediated innate immune response in aged mice fed alcohol. Notably, recent studies have highlighted a dark side of persistent TFEB activation, which may lead to cell fate changes, polycystic liver disease, and even liver tumorigenesis (Pastore, Huynh et al. 2020). Therefore, more studies are needed to evaluate the long-term effects of autophagy and TFEB activation in aging ALD.

In conclusion, this study highlights the role of advanced age in perpetuating the negative impact of heavy alcohol consumption on key hepatic processes, including lipid and fatty acid metabolism, inflammatory signaling, and TFEB-mediated autophagy. Additionally, alcohol-influenced induction of cellular senescence remains persistent in old age, potentially impacting the overall health and function of the aged liver. Activation of hepatic TFEB is an effective mechanism to counter age- and ethanol-induced upregulation of pro-inflammatory immune signaling.

## Acknowledgement

This study was partly supported by the National Institute of Health (NIH) funds R37 AA020518, R01 AG072895 and R01AA031230 (WXD).

## Abbreviations

AD: Adenovirus
ADH: Alcohol dehydrogenase
AH: Alcohol-associated hepatitis
ALD: Alcohol-associated liver disease
ALDH2: Acetylaldehyde dehydrogenase 2
ALT: Alanine transaminase
ASH: Alcohol-induced steatohepatitis
AST: Aspartate aminotransferase
ATP6V1A1: ATPase *H^+^* transporting V1 subunit A1
ATP6V1B2: ATPase *H^+^* transporting V1 subunit B2
BHBA: Betahydroxybutyrate
CCL6: Chemokine ligand 6
CD36: Cluster of differentiation 36
cGAS: Cyclic GMP-AMP synthase
CPT1α: Carnitine palmitoyltransferase 1A
CYP2E1: Cytochrome P450 2E1
DAGs: Diacylglycerols
DNL: *De novo* lipogenesis
EtOH: Ethanol
ER: Endoplasmic Reticulum
FA: Fatty acid
FAO: Fatty acid oxidation
IL-8: Interleukin-8
IRF-3: Interferon-3
IRF-7: Interferon-7
IGFBP-1: Insulin growth factor-binding protein-1
KCs: Kupffer cells
LAMP-1: Lysosome associated membrane protein-1
LC3-II: Microtubule-associated protein 1A/1B-light chain 3
LDL-R: Low-density lipoprotein-receptor
MAGs: Monoacylglycerols
MCP-1: Monocyte chemoattractant protein-1
NF-κB: Nuclear factor kappa-light-chain-enhancer of activated B cells
p62: Sequestosome-1
PCA: Principle component analysis
ROS: Reactive oxygen species
RT-qPCR: Real-time quantitative polymerase chain reaction
SA β-gal: Senescence-associated beta-galactosidase
SAMHSA: Substance Abuse and Mental Health Services Administration
SASP: Senescence-associated secretory phenotype
STING: stimulator of interferon genes
TFEB: Transcription factor EB
TGs: Triglycerides
TNFα: Tumor necrosis factor alpha

## Notes

### Competing Interest Statement

The authors have declared no competing interest.

## References

Abokyi, S., G. Ghartey-Kwansah and D. Y. Y. Tse (2023). “TFEB is a central regulator of the aging process and age-related diseases.” Ageing Research Reviews 89.

Ajoolabady, A., D. Pratico, S. Bahijri, B. Eldakhakhny, J. Tuomilehto, F. Wu and J. Ren (2025). “Hallmarks and mechanisms of cellular senescence in aging and disease.” Cell Death Discov 11(1): 364.

Aravinthan, A., G. Pietrosi, M. Hoare, J. Jupp, A. Marshall, C. Verrill, S. Davies, A. Bateman, N. Sheron, M. Allison and G. J. Alexander (2013). “Hepatocyte expression of the senescence marker p21 is linked to fibrosis and an adverse liver-related outcome in alcohol-related liver disease.” PLoS One 8(9): e72904.

Baechle, J. J., N. Chen, P. Makhijani, S. Winer, D. Furman and D. A. Winer (2023). “Chronic inflammation and the hallmarks of aging.” Mol Metab 74: 101755.

Bertola, A., S. Mathews, S. H. Ki, H. Wang and B. Gao (2013). “Mouse model of chronic and binge ethanol feeding (the NIAAA model).” Nat Protoc 8(3): 627–637.

Cao, P., X. Chao, H. M. Ni and W. X. Ding (2026). “An Update on Animal Models of Alcohol-Associated Liver Disease.” Am J Pathol 196(1): 4–19.

Ceni, E., T. Mello and A. Galli (2014). “Pathogenesis of alcoholic liver disease: role of oxidative metabolism.” World J Gastroenterol 20(47): 17756–17772.

Chao, X., S. Wang, K. Zhao, Y. Li, J. A. Williams, T. Li, H. Chavan, P. Krishnamurthy, X. C. He, L. Li, A. Ballabio, H. M. Ni and W. X. Ding (2018). “Impaired TFEB-Mediated Lysosome Biogenesis and Autophagy Promote Chronic Ethanol-Induced Liver Injury and Steatosis in Mice.” Gastroenterology 155(3): 865–879 e812.

Chao, X. J., S. G. Wang, X. W. Ma, C. Zhang, H. Qian, S. N. Williams, Z. L. Sun, Z. Y. Peng, W. Q. Liu, F. Li, N. Sheshadri, W. X. Zong, H. M. Ni and W. X. Ding (2023). “Persistent mTORC1 activation due to loss of liver tuberous sclerosis complex 1 promotes liver injury in alcoholic hepatitis.” Hepatology 78(2): 503–517.

Chen, H., K. Hinz, C. Zhang, Y. Rodriguez, S. N. Williams, M. Niu, X. Ma, X. Chao, A. L. Frazier, K. E. McCarson, X. Wang, Z. Peng, W. Liu, H. M. Ni, J. Zhang, R. H. Swerdlow and W. X. Ding (2024). “Late-Life Alcohol Exposure Does Not Exacerbate Age-Dependent Reductions in Mouse Spatial Memory and Brain TFEB Activity.” Biomolecules 14(12).

Dukic, M., T. Radonjic, I. Jovanovic, M. Zdravkovic, Z. Todorovic, N. Kraisnik, B. Arandelovic, O. Mandic, V. Popadic, N. Nikolic, S. Klasnja, A. Manojlovic, A. Divac, J. Gacic, M. Brajkovic, S. Opric, M. Popovic and M. Brankovic (2023). “Alcohol, Inflammation, and Microbiota in Alcoholic Liver Disease.” Int J Mol Sci 24(4).

Hinz, K., H. Qian, B. Peiffer, Z. L. Sun, H. M. Ni and W. X. Ding (2025). “Role of SQSTM1/p62 in regulating Mallory-Denk body in alcohol-associated liver disease.” Egastroenterology 3(4).

Hong, X., S. Huang, H. Jiang, Q. Ma, J. Qiu, Q. Luo, C. Cao, Y. Xu, F. Chen, Y. Chen, C. Sun, H. Fu, Y. Liu, C. Li, F. Chen and P. Qiu (2024). “Alcohol-related liver disease (ALD): current perspectives on pathogenesis, therapeutic strategies, and animal models.” Front Pharmacol 15: 1432480.

Hunt, N. J., S. W. S. Kang, G. P. Lockwood, D. G. Le Couteur and V. C. Cogger (2019). “Hallmarks of Aging in the Liver.” Comput Struct Biotechnol J 17: 1151–1161.

Jeon, S. and R. Carr (2020). “Alcohol effects on hepatic lipid metabolism.” J Lipid Res 61(4): 470–479.

Julien, J., T. Ayer, E. B. Tapper and J. Chhatwal (2024). “The Rising Costs of Alcohol-Associated Liver Disease in the United States.” Am J Gastroenterol 119(2): 270–277.

Kim, I. H., T. Kisseleva and D. A. Brenner (2015). “Aging and liver disease.” Curr Opin Gastroenterol 31(3): 184–191.

Kitada, M. and D. Koya (2021). “Autophagy in metabolic disease and ageing.” Nature Reviews Endocrinology 17(11): 647–661.

Le Couteur, D. G., M. C. Ngu, N. J. Hunt, A. E. Brandon, S. J. Simpson and V. C. Cogger (2025). “Liver, ageing and disease.” Nature Reviews Gastroenterology & Hepatology 22(10): 680–695.

Lopez-Otin, C., M. A. Blasco, L. Partridge, M. Serrano and G. Kroemer (2023). “Hallmarks of aging: An expanding universe.” Cell 186(2): 243–278.

Ma, X., A. Chen, L. Melo, A. Clemente-Sanchez, X. Chao, A. R. Ahmadi, B. Peiffer, Z. Sun, H. Sesaki, T. Li, X. Wang, W. Liu, R. Bataller, H. M. Ni and W. X. Ding (2023). “Loss of hepatic DRP1 exacerbates alcoholic hepatitis by inducing megamitochondria and mitochondrial maladaptation.” Hepatology 77(1): 159–175.

Ma, X., H. M. Ni and W. X. Ding (2024). “Perspectives of mitochondria-lysosome-related organelle in hepatocyte dedifferentiation and implications in chronic liver disease.” eGastroenterology 2(1).

Ma, X. W., M. W. Niu, H. M. Ni and W. X. Ding (2026). “Mitochondrial dynamics, quality control, and mtDNA in alcohol-associated liver disease and liver cancer.” Hepatology 83(3): 639–660.

Mackowiak, B., Y. Fu, L. Maccioni and B. Gao (2024). “Alcohol-associated liver disease.” J Clin Invest 134(3).

Mandrekar, P. and A. Mandal (2024). “Pathogenesis of Alcohol-Associated Liver Disease.” Clin Liver Dis 28(4): 647–661.

Massey, V. L. and G. E. Arteel (2012). “Acute alcohol-induced liver injury.” Front Physiol 3: 193.

Nagy, L. E., W. X. Ding, G. Cresci, P. Saikia and V. H. Shah (2016). “Linking Pathogenic Mechanisms of Alcoholic Liver Disease With Clinical Phenotypes.” Gastroenterology 150(8): 1756–1768.

Nakamura, J., T. Yamamoto, Y. Takabatake, T. Namba-Hamano, A. Takahashi, J. Matsuda, S. Minami, S. Sakai, H. Yonishi, S. Maeda, S. Matsui, H. Kawai, I. Matsui, T. Yamamuro, R. Edahiro, S. Takashima, A. Takasawa, Y. Okada, T. Yoshimori, A. Ballabio and Y. Isaka (2024). “Age-related TFEB downregulation in proximal tubules causes systemic metabolic disorders and occasional apolipoprotein A4-related amyloidosis.” JCI Insight 10(3).

Narro, G. E. C., L. A. Diaz, E. K. Ortega, M. F. B. Garin, E. C. Reyes, P. S. M. Delfin, J. P. Arab and R. Bataller (2024). “Alcohol-related liver disease: A global perspective.” Ann Hepatol 29(5): 101499.

Nguyen, P., V. Leray, M. Diez, S. Serisier, J. Le Bloc’h, B. Siliart and H. Dumon (2008). “Liver lipid metabolism.” J Anim Physiol Anim Nutr (Berl) 92(3): 272–283.

Niu, M., X. Wei, X. Ma and W. X. Ding (2026). “Megamitochondria in alcohol-associated liver disease and cancer: a friend or foe?” eGastroenterology 4(2): e100408.

Pastore, N., T. Huynh, N. J. Herz, A. Calcagni, T. J. Klisch, L. Brunetti, K. H. Kim, M. De Giorgi, A. Hurley, A. Carissimo, M. Mutarelli, N. Aleksieva, L. D’Orsi, W. R. Lagor, D. D. Moore, C. Settembre, M. J. Finegold, S. J. Forbes and A. Ballabio (2020). “TFEB regulates murine liver cell fate during development and regeneration.” Nat Commun 11(1): 2461.

Pyo, J. O., S. M. Yoo, H. H. Ahn, J. Nah, S. H. Hong, T. I. Kam, S. Jung and Y. K. Jung (2013). “Overexpression of Atg5 in mice activates autophagy and extends lifespan.” Nature Communications 4.

Qian, H., X. Chao, S. Wang, Y. Li, X. Jiang, Z. Sun, T. Rulicke, K. Zatloukal, H. M. Ni and W. X. Ding (2023). “Loss of SQSTM1/p62 Induces Obesity and Exacerbates Alcohol-induced Liver Injury in Aged Mice.” Cell Mol Gastroenterol Hepatol.

Qian, H., X. Chao, J. Williams, S. Fulte, T. Li, L. Yang and W. X. Ding (2021). “Autophagy in liver diseases: A review.” Mol Aspects Med: 100973.

Tian, T., Y. Xue, Z. Song, B. Jin-Smith, J. Barkin, M. Ottallah, M. Mannan, A. Zhirkova, D. Zhou and L. Pi (2025). “Targeted clearance of senescent cells alleviates alcohol-associated liver disease by restoring cellular function and immune balance.” Geroscience.

Timchenko, N. A. (2009). “Aging and liver regeneration.” Trends Endocrinol Metab 20(4): 171–176.

Uddin, M. N., N. Nishio, S. Ito, H. Suzuki and K. Isobe (2012). “Autophagic activity in thymus and liver during aging.” Age (Dordr) 34(1): 75–85.

Ueno, T. and M. Komatsu (2017). “Autophagy in the liver: functions in health and disease.” Nature Reviews Gastroenterology & Hepatology 14(3): 170–184.

Uyar, B., D. Palmer, A. Kowald, H. Murua Escobar, I. Barrantes, S. Moller, A. Akalin and G. Fuellen (2020). “Single-cell analyses of aging, inflammation and senescence.” Ageing Res Rev 64: 101156.

Wan, Y., K. McDaniel, N. Wu, S. Ramos-Lorenzo, T. Glaser, J. Venter, H. Francis, L. Kennedy, K. Sato, T. Zhou, K. Kyritsi, Q. Huang, T. Annable, C. Wu, S. Glaser, G. Alpini and F. Meng (2017). “Regulation of Cellular Senescence by miR-34a in Alcoholic Liver Injury.” Am J Pathol 187(12): 2788–2798.

Wang, H., M. K. Muthu Karuppan, D. Devadoss, M. Nair, H. S. Chand and M. K. Lakshmana (2021). “TFEB protein expression is reduced in aged brains and its overexpression mitigates senescence-associated biomarkers and memory deficits in mice.” Neurobiol Aging 106: 26–36.

Wang, Y. X. J. (2021). “Gender-specific liver aging and magnetic resonance imaging.” Quant Imaging Med Surg 11(7): 2893–2904.

Williams, J. A., S. Manley and W. X. Ding (2014). “New advances in molecular mechanisms and emerging therapeutic targets in alcoholic liver diseases.” World J Gastroenterol 20(36): 12908–12933.

Williams, S. N. and W. X. Ding (2025). “The impact of aging on liver health and the development of liver diseases.” Hepatology Communications 9(10).

Williams, S. N. and W. X. Ding (2025). “The impact of aging on liver health and the development of liver diseases.” Hepatol Commun 9(10).

Xu, H. and H. Wang (2022). “Immune cells in alcohol-related liver disease.” Liver Res 6(1): 1–9.

